# Genetic dissection of BDNF and TrkB expression in glial cells

**DOI:** 10.1101/2023.07.14.549007

**Authors:** Changran Niu, Xinpei Yue, Juan Ji An, Haifei Xu, Baoji Xu

**Affiliations:** Department of Neuroscience, The Herbert Wertheim UF Scripps Institute for Biomedical Innovation & Technology, University of Florida, Jupiter, 33458, FL; Skaggs Graduate School of Chemical and Biological Sciences, The Scripps Research Institute, La Jolla, 92037, CA

**Keywords:** BDNF, TrkB, Cre, glia, microglia, astrocyte, oligodendrocyte

## Abstract

The brain-derived neurotrophic factor (BDNF) and its high-affinity receptor tropomyosin-related kinase receptor B (TrkB) are widely expressed in the central nervous system. It is well documented that neurons express BDNF and full-length TrkB (TrkB.FL), and a lower level of truncated TrkB (TrkB.T). With conflicting results, glial cells also have been reported to express BDNF and TrkB. In the current study, we employed a more sensitive and reliable genetic method to characterize the expression of BDNF and TrkB in glial cells in the mouse brain. We utilized three Cre mouse strains in which Cre recombinase is expressed in the same cells as BDNF, TrkB.FL, or all TrkB isoforms, and crossed them to Cre-dependent EGFP reporter mice to label BDNF- or TrkB-expressing cells. We performed immunohistochemistry with glial cell markers to examine the expression of BDNF and TrkB in microglia, astrocytes, and oligodendrocytes. Surprisingly, we found no BDNF- or TrkB-expressing microglia in the brain and spinal cord. Consistent with previous studies, most astrocytes only express TrkB.T in the adult brain. Moreover, there are a small number of astrocytes and oligodendrocytes that express BDNF, the function of which is to be determined. We also found that oligodendrocyte precursor cells, but not mature oligodendrocytes, express both TrkB.FL and TrkB.T in the adult brain. These results not only clarify the expression of BDNF and TrkB in glial cells, but also open opportunities to investigate previously unidentified roles of BDNF and TrkB in glial cells.

## 1. Introduction

Brain-derived neurotrophic factor (BDNF) is a neurotrophin that is widely expressed in the central nervous system. BDNF signals via the tropomyosin-related kinase B (TrkB) receptor to regulate neuronal proliferation, differentiation and survival, and synaptic function [1]. Binding of BDNF to the full-length TrkB (TrkB.FL) receptor triggers dimerization of TrkB.FL and tyrosine autophosphorylation, leading to activation of three main downstream signaling cascades mediated by phospholipase C γ1 (PLCγ1), phosphoinositide 3-kinase, and RAS/RAF/mitogen-activated protein kinase, respectively [2]. Alternative splicing of *Ntrk2* exons produces the truncated TrkB (TrkB.T) isoforms that lack nearly all the intracellular domain including the tyrosine kinase motif [3]. TrkB.T can negatively regulate TrkB.FL signaling by forming a TrkB.FL-TrkB.T heterodimer and inhibiting the activation of TrkB.FL [4]. TrkB.T also functions to sequester and translocate BDNF, induce neurite outgrowth, and activate downstream protein kinase C and PLCγ via G-protein and inhibit Rho GTPase via dissociating Rho GDP dissociation inhibitor [4]. Because of the diverse roles of the BDNF-TrkB pathway, dysregulation of the BDNF-TrkB signaling has been implicated in various disease conditions, including cognitive impairment, obesity, neurodegenerative diseases, and cancer [5].

BDNF-TrkB signaling pathway has been studied extensively in neurons. It is well-documented that neurons express BDNF, TrkB.FL and a lower level of TrkB.T. Expression of BDNF and TrkB has also been reported in glial cells. BDNF expressed in microglia was suggested to be involved in neuropathic pain and motor-learning dependent synapse formation [6, 7]. However, other studies showed very low or no expression of BDNF in microglia [8]. Therefore, it is debatable whether microglia express BDNF or not. It has been shown that astrocytes predominantly express TrkB.T [1, 9]. Whether astrocytes also express TrkB.FL is not clear yet. Some studies have shown TrkB.FL expression in astrocytes isolated from mouse brain [10, 11], whereas other studies show no TrkB.FL expression in astrocytes [12, 13]. Astrocytes can internalize proBDNF secreted by neurons, convert proBDNF to mature BDNF (mBDNF) and release mBDNF for synaptic reuse [14, 15]. Astrocytes was shown to express BDNF *in vitro* which regulates neuronal dendrite maturation [16], but whether astrocytes express BDNF *in vivo* under physiological conditions is unknown. Oligodendrocytes have also been shown to express BDNF, the reduction of which impairs presynaptic vesicular exocytosis [17]. Oligodendroglial expression of TrkB is important for oligodendrocyte myelination and oligodendrocyte precursor cell (OPC) proliferation [18–20]. But it has not been characterized which TrkB isoform oligodendrocytes express and what percentage of oligodendrocytes express BDNF or TrkB.

To fill these knowledge gaps, in the current study, we employed a genetic method to label and characterize the expression of BDNF and TrkB in glial cells, including microglia, astrocytes and oligodendrocytes. We crossed the *Bdnf*^2A-Cre/+^ [21], *Ntrk2*^2A-Cre/+^, and *Ntrk2*^CreER/+^ [22] mouse strains to the Cre-dependent B6;129S4-Gt(*ROSA*)*26Sor^tm9(EGFP/Rpl10a)Amc^*/J (EGFP-L10a) reporter mice [23] to generate double heterozygous mice in which cells expressing BDNF or TrkB were labeled by EGFP. We then performed immunofluorescence staining of cell type markers with the brain sections from these mice. Colocalization of EGFP and cell type markers was analyzed to determine the expression of BDNF and TrkB in glial cells. There are several advantages of this method: (1) high sensitivity because a very small amount of Cre expression can induce EGFP expression in the reporter mice; (2) no involvement of BDNF and TrkB antibodies which have non-specific immunoreactivity; (3) no need to isolate glia from the mouse brain, which involves potential contamination of other cell types; (4) being able to observe expression of BDNF and TrkB *in vivo* at the level of single cells.

Our results show that homeostatic and activated microglia do not express either BDNF or TrkB. Astrocytes predominantly express TrkB.T, and a small astrocyte population express TrkB.FL and BDNF in the adult mice. A subset of oligodendrocytes express BDNF in the hippocampal CA1, as well as TrkB.T and TrkB.FL.

## 2. Materials and Methods

### 2.1 Animals

Mice of the EGFP-L10a/+ (B6;129S4-Gt(*ROSA*)*26Sor^tm9(EGFP/Rpl10a)Amc^*/J; stock #024750) and the *Bdnf*^2A-Cre/+^ (B6.FVB-*Bdnf^em1)cre_Zak^*/J; stock #030189) strains were obtained from the Jackson Laboratory. The *Ntrk2*^CreER/+^ mouse strain was generously provided by Dr. David Ginty at Harvard Medical School [22]. *Ntrk2*^2A-Cre/+^, *Ntrk2*^CreER/+^, and *Bdnf*^2A-Cre/+^mice were crossed to EGFP-L10a/+ mice to generate *Ntrk2*^2A-Cre/+^;EGFP-L10a/+, *Ntrk2*^CreER/+^;EGFP-L10a/+ and *Bdnf*^2A-Cre/+^;EGFPL10a/+ mice. All experiments were performed in accordance with relevant guidelines and regulations regarding the use of experimental animals. The Animal Care and Use Committees at the UF Scripps approved all animal procedures used in this study.

### 2.2 Generation of *Ntrk2*^2A-Cre/+^ mice

We generated a *Ntrk2^2A-Cre^* mouse allele in which the DNA sequence encoding P2A-Cre recombinase was inserted immediately before the stop codon for TrkB.FL at the *Ntrk2* locus using the CRISPR/Cas9 technique. sgRNA (5’-AGTCAAGAGGTTCGTCGTGT-3’) was designed using the CRISPR tool (http://crispr.mit.edu) to minimize potential off-target effects. The DNA donor plasmid contains two homology arms with 840-bp flanking the P2A-Cre sequence, which were cloned into BamHI digested pBluescript II KS (-) vector using the Gibson assembly method (NEB, E5510). To block further Cas9 targeting and recutting after undergoing homology-direct repair, we introduced a silent mutation into the PAM motif of the sgRNA located within the 3’ homology arm in the donor plasmid by using Q5 Site-Directed Mutagenesis kit (NEB, E0054). The donor plasmid was confirmed with DNA sequencing. Microinjection of a mixture of sgRNA, donor DNA, and Cas9 protein was performed by the Genomic Modification Facility at Scripps Research. Zygotes were cultured to the blastocyst stage *in vitro*. Genomic DNA from blastocysts was extracted and PCR screen was performed to select blastocysts with successful homologous recombination. Positive blastocysts were transferred into the oviduct of pseudo-pregnant females to produce founder mice. Two out of twenty-eight pups were positive for the knockin, confirmed by the genomic DNA PCR using the following primers: CCTCCTGGTGAGCAAACGAT (forward) and GACATAGGGCCGGGATTCTC (reverse) for PCR across the 5’ homology arm, and CACCTCCATGCCTGTGTTTT (forward) and GGTTCTTGCGAACCTCATCA (reverse) for PCR across the 3’ homology arm. The founder mice were crossed to C57BL/6J to produce mice with germline transmission. Cre expression in these lines were confirmed by *in situ* hybridization.

### 2.3 In situ hybridization

Freshly dissected mouse brains were quickly frozen in a 2-methylbutane (Fisher Scientific #O3551-4) dry ice bath. Subsequently, 14-μm thick cryostat sections were collected onto Superfrost Plus slides (Fisherbrand #12-550-15) and used for in situ hybridization. Fluorescence in situ hybridization was performed using the RNAscope Multiplex Fluorescent Reagent Kit v2 (#323100, Advanced Cell diagnostics). Brain sections were fixed with pre-chilled 10% formalin at 4°C for 15 min, and dehydrated with 50%, 75% and 100% ethanol. Sections were air-dried for 20 min and followed by v2Hydrogen peroxide for 10 min and pretreated with protease IV for 30 min, washed 2 times in PBS and incubated with the target probes (Ntrk2-C3, #423611-C3; Cre-C2, #31228-C2) for 2 hours at 40°C. Signals were amplified by subsequent incubation with v2Amp1 for 30 min, v2Amp2 for 30 min, v2Amp3 for 15 min, v2C2/C3HPR for 15 min and followed by TSA Plus Fluorescein/Cyanine5 (NEL754001KT, Akoya Biosciences) for 30 min at 40°C. Slides were counterstained with DAPI, and images were acquired using a Nikon C2+ confocal microscope.

### 2.4 Immunohistochemistry

Mice were perfused with phosphate-buffered saline (PBS) and then 4% paraformaldehyde (PFA) in PBS. The brain was dissected and postfixed overnight in 4% PFA-PBS. Mouse brains were soaked in 30% sucrose-PBS for cryoprotection. Brains were sectioned into 40-μm-thick slices using a sliding microtome (Leica). Brain sections were blocked in 0.3% Triton X-100 and 10% normal horse serum before overnight incubation in primary antibody at 4 °C. The following primary antibodies were used: guinea pig anti-Iba1(1:500; Synaptic System #234308), goat anti-SOX9 (1:2000; R&D Systems #AF3075), mouse anti-GFAP (1:500; Cell Signaling Technology #3670S), mouse anti-Olig2 (1:250; Millipore #MABN50), rabbit anti-ASPA (1:1500; GeneTex #GTX113389), chicken anti-GFP (1:2000; abcam #13970), rabbit anti-DsRed (1:1000; TaKaRa #632496). The brain sections were washed three times with PBS before incubation in secondary antibody for 1 hour at room temperature. The brain sections were then counterstained with DAPI, mounted onto microscope slides, and covered with mounting media.

### 2.5 LPS injection

LPS (Millipore #L6529) was dissolved in sterile saline to a final concentration of 0.1 μg/μL. The mice were given daily saline or LPS injection at a dose of 500 μg/kg body weight for seven consecutive days. The mice were harvested the next day after the last treatment.

### 2.6 Tamoxifen injection

Tamoxifen (Sigma #T5648-5G) was dissolved in 100% ethanol to a concentration of 20 mg/ml. An equal volume of corn oil was added to the tamoxifen solution. The corn oil/ethanol mixture was vortexed, centrifuged for 10 mins at 13,000 rpm, and then placed in a vacuum centrifuge for 30 mins to evaporate ethanol. Tamoxifen was given to *Ntrk2*^CreER/+^;EGFP-L10a/+ mice at the age of 6 weeks through daily i.p. injection at a dose of 150 mg/kg body weight for 10 consecutive days to induce translocation of the Cre-ER fusion protein to the nucleus. The mice were given three weeks to recover before they were harvested.

### 2.7 Plasmids and viruses

pAAV8-EF1a-Nuc-flox(mCherry)-EGFP was obtained from Addgene (plasmid #112677, a gift from Brandon Harvey [24]). For higher expression in glial cells, the EF1a promoter was replaced by the CAG promoter from pAAV-CAG-mNeonGreen (Addgene plasmid #99134, a gift from Viviana Gradinaru [25]). pAAV8-EF1a-Nuc-flox(mCherry)-EGFP was digested with AgeI and BamHI, and the CAG promoter fragment was amplified by PCR (forward primer: GAAT ACCGGTCGTTACATAACTTACGGTAAATG; reverse primer: GAATGGATCCCGCCCGCCGCGCGCTT). PCR product was digested with AgeI and BamHI and ligated into the backbone of pAAV8-EF1a-Nuc-flox(mCherry)-EGFP. The resulting pAAV8-CAG-Nuc-flox(mCherry)-EGFP plasmid was confirmed by sequencing and used to package AAV viruses. AAV viruses were purified with TaKaRa AAVpro Purification Kit Maxi (#6666).

### 2.8 Stereotaxic injection of adeno-associated virus

Mice were anesthetized using isoflurane and securely positioned on a stereotaxic holder (David Kopf, Model 940). A small incision was made to expose the skull, and subsequently a small hole was drilled on the skull above an injection site. For injection, a Nanofil 33-gauge needle (World Precision Instruments, #NF33BV-2) was slowly inserted into the CA1 region (AP: -1.60 mm; ML: ± 0.80 mm; DV: −1.80 mm). AAV8-CAG-Nuc-flox(mCherry)-EGFP (150 nl at 3.12E+12 vg/mL) was then injected at a rate of 30 nl/min using a micro syringe pump (World Precision Instruments, SYS-MICRO4). The needle was then withdrawn after staying in the same position for 5 min. Following the injection, mice received Loxicom (5 mg/kg) for analgesia and were returned to their home cages.

### 2.9 Imaging and Cell number quantification

Fluorescent images were captured using a Nikon C2+ confocal microscope. Cell numbers were quantified using the cell counter plugin in FIJI software.

## 3. Results

### 3.1 Homeostatic and activated microglia in the cortex, hippocampus and spinal cord do not express BDNF

*Bdnf*^2A-Cre/+^ mice that express Cre recombinase in BDNF-expressing cells were crossed to Cre-dependent EGFPL10a/+ reporter mice to generate *Bdnf*^2A-Cre/+^;EGFPL10a/+ mice. Cre-mediated excision of *loxP*-flanked STOP sequence results in constitutive expression of EGFP localized in the nucleus and cytoplasm of Cre-expressing cells. EGFP is thus expressed in cells that express BDNF at any time during the lifespan of a *Bdnf*^2A-Cre/+^;EGFP-L10a/+ mouse. We first used NeuN as a marker to label neurons. As expected, most neurons in the cortex and hippocampal CA1 express BDNF (Fig. 1A-F).

**Figure 1.**
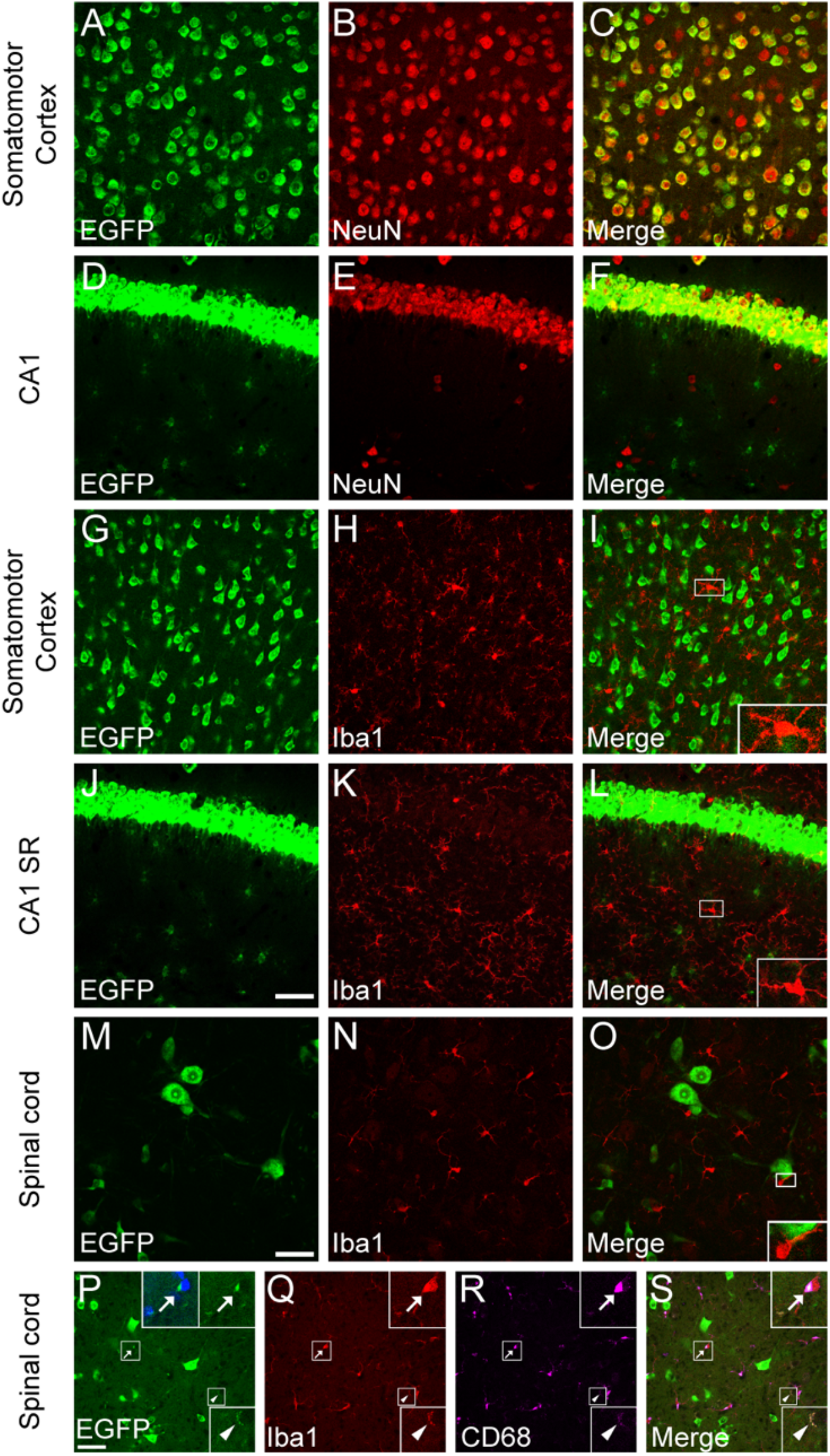
Homeostatic microglia do not express BDNF. (**A-F**) BDNF expression in NeuN^+^ neurons in the MO (A-C) and CA1 (D-F). (**G-O**) Homeostatic microglia do not express BDNF in the MO (G-I), CA1 (J-L), and spinal cord (M-O). (**P-S**) Colocalization of EGFP puncta signals with CD68^+^ microglia lysosomes. Arrows denote EGFP signal inside microglial cytoplasm. Arrowheads denote EGFP signal inside microglial processes. Scale bar = 50 μm. n = 2 mice.

Next, we wanted to know whether microglia express BDNF. For this, we used Iba1 antibody to label microglia in *Bdnf*^2A-Cre/+^;EGFP-L10a/+ mice. Surprisingly, we did not observe EGFP expression in nuclei of Iba1^+^ microglia in the somatomotor cortex (MO), hippocampal CA1 stratum radiatum (SR), or the spinal cord of 4-month-old *Bdnf*^2A-Cre/+^;EGFPL10a/+ mice, indicating that homeostatic microglia do not express BDNF under physiological conditions (Fig. 1G-O). However, we did observe EGFP puncta in some microglial cell bodies and processes, outside microglial nuclei (Fig. 1P-S). Immunostaining of CD68, a microglial lysosomal marker, showed that the EGFP puncta are always colocalized with CD68 (Fig. 1P-S). This indicates that these EGFP signals in microglia come from BDNF-expressing cells engulfed by microglia and are not from BDNF expression in microglia.

Microglia can exist in homeostatic or activated states. To investigate whether activated microglia express BDNF under pathological conditions, we injected lipopolysaccharide (LPS) to induce microglial activation in adult *Bdnf*^2A-Cre/+^;EGFPL10a/+ mice. CD68 is used as a marker for microglia activation, as activated microglia express a higher level of CD68. LPS treatment resulted in a significant increase in CD68 intensity per microglia (Fig. 2). We did not observe EGFP signal in Iba1^+^ microglial nuclei in LPS-injected mice, suggesting that activated microglia also do not express BDNF (Fig. 2E-H). Therefore, our results suggest that both homeostatic and activated microglia do not express BDNF.

**Figure 2.**
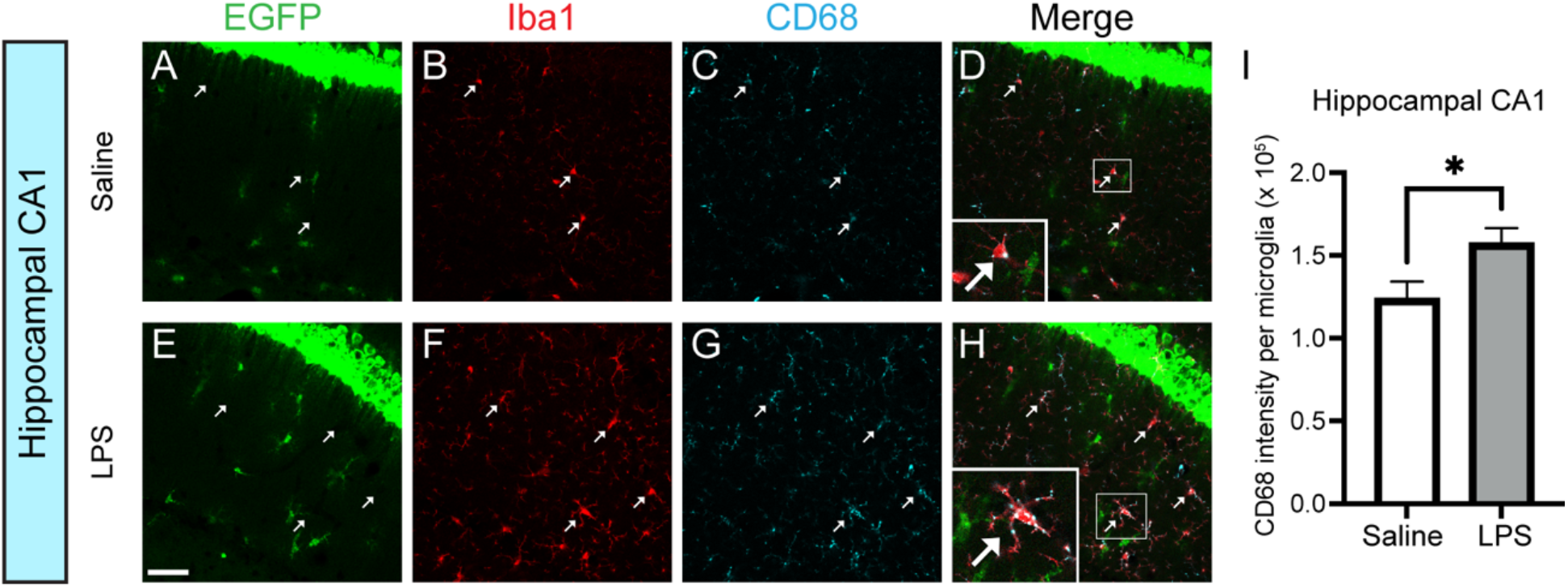
Activated microglia do not express BDNF. (**A-H**) Immunostaining of Iba1 and CD68 in the hippocampus of *Bdnf*^2A-Cre/+^;EGFPL10a/+ mice injected with saline (A-D) or LPS (E-H). Arrows indicate colocalization of Iba1 and CD68, and no EGFP inside microglia. Scale bar = 50 μm. (**I**) Quantification of CD68 intensity per microglia. n = 1 35 microglia from one saline mouse and 50 microglia from 2 LPS mice. *p = 0.0204 by two-sided *t*-test. Data are shown as mean ± s.e.m.

### 3.2 Some astrocytes express BDNF in the cortex and hippocampus

Whether astrocytes express BDNF *in vivo* under physiological conditions is unknown. To answer this question, we labeled astrocytes in *Bdnf*^2A-Cre/+^;EGFPL10a/+ mice using SOX9 antibody for the somatosensory cortex (SS) and GFAP antibody for the hippocampus. We observed that a small portion of astrocytes (2.6% in the SS and 18.3% in the CA1 SR) are EGFP^+^ in the *Bdnf*^2A-Cre/+^;EGFPL10a/+ mice, indicating that these cells express BDNF at some point during the lifetime of the mice (Fig. 3A-H). The EGFP signals in these mice are cumulative, because once the EGFP expression in a cell is turned on by Cre, it is never turned off even if the cell stops to express Cre. Therefore, we cannot know when BDNF is expressed in these mice.

**Figure 3.**
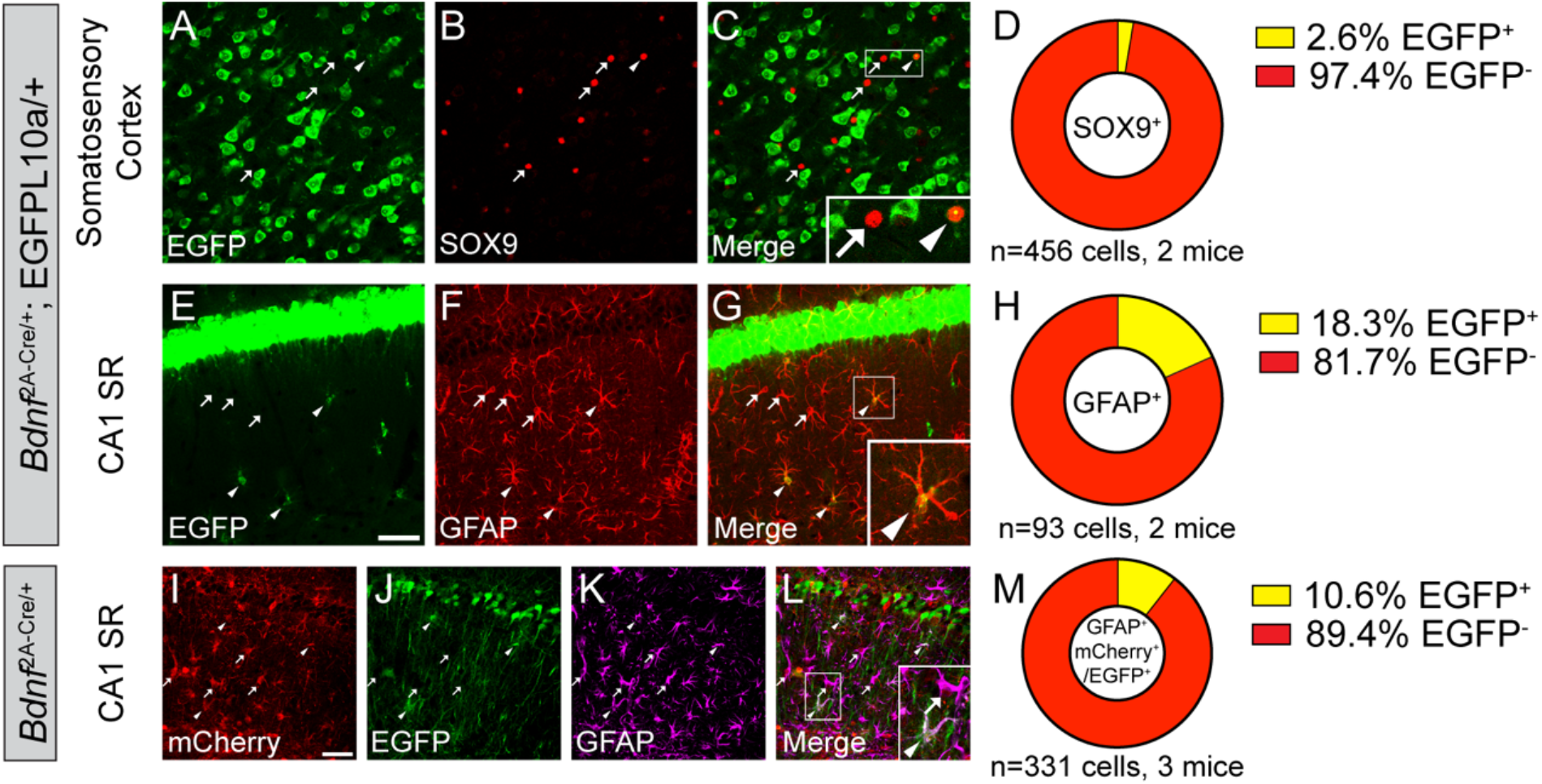
A small subset of astrocytes express BDNF in the hippocampus. (**A-H**) Immunostaining of SOX9 (A-C) and GFAP (E-G) in the SS (A-C) and CA1 SR (E-G) of *Bdnf*^2A-Cre/+^;EGFPL10a/+ mice, and (D, H) quantification of colocalization. (**I-M**) Immunostaining of GFAP in the CA1 SR of adult *Bdnf*^2A-Cre/+^ mice injected with the mCherry-to-EGFP color switch AAV, and (M) quantification of colocalization. Scale bar = 50 μm. Arrowheads denote EGFP-expressing astrocytes. Arrows denote astrocytes that do not express EGFP.

To test whether astrocytes in adult mice still express BDNF, we injected a Cre-dependent mCherry-to-EGFP color switch adeno-associated virus (AAV), AAV8-CAG-Nuc-flox(mCherry)-EGFP, into the hippocampal CA1 of 2-month-old *Bdnf*^2A-Cre/+^ mice. In these mice, transduced Cre-negative cells will express mCherry, while transduced Cre-positive cells will express EGFP. Around 87% astrocytes were transduced by the AAV, and 10.6% of the transduced astrocytes express BDNF in adult mice (Fig. 3I-M). This suggests that a small portion of astrocytes in the CA1 SR still express BDNF in adult mice. Because this number is smaller than the cumulative 18.3% in *Bdnf*^2A-Cre/+^;EGFPL10a/+ mice, there could be two possibilities: (1) some astrocytes only express BDNF early during development and stop BDNF expression in adults; (2) at each different time point, there is a different small subset of astrocytes that express BDNF, and that accumulates to the 18.3% seen in *Bdnf*^2A-Cre/+^;EGFPL10a/+ mice. To figure out which possibility is true, future studies can investigate BDNF expression at developmental stages.

### 3.3 A small portion of oligodendrocytes express BDNF in the hippocampal CA1 SR

Next, we examined BDNF expression in Olig2^+^ oligodendrocytes. We found that most Olig2^+^ oligodendrocytes in the SS do not express BDNF, and 4.4% of Olig2^+^ oligodendrocytes in the CA1 SR express BDNF at some point during the lifespan of *Bdnf*^2A-Cre/+^;EGFPL10a/+ mice (Fig. 4A-H). To find out whether these oligodendrocytes express BDNF in adult mice, we injected the mCherry-to-EGFP color switch virus to the hippocampus of *Bdnf*^2A-Cre/+^ mice and examined colocalization of Olig2 with mCherry and EGFP. Out of the transduced Olig2^+^ cells, 14.5% express BDNF in adult mice (Fig. 4I-M). It is unexpected that the percentage of BDNF-expressing oligodendrocytes in adult is higher than that of cumulative expression in *Bdnf*^2A-Cre/+^;EGFPL10a/+ mice. One possible explanation is that AAV injection might cause tissue damage or myelin damage, which induced the increase in BDNF expression in oligodendrocytes.

**Figure 4.**
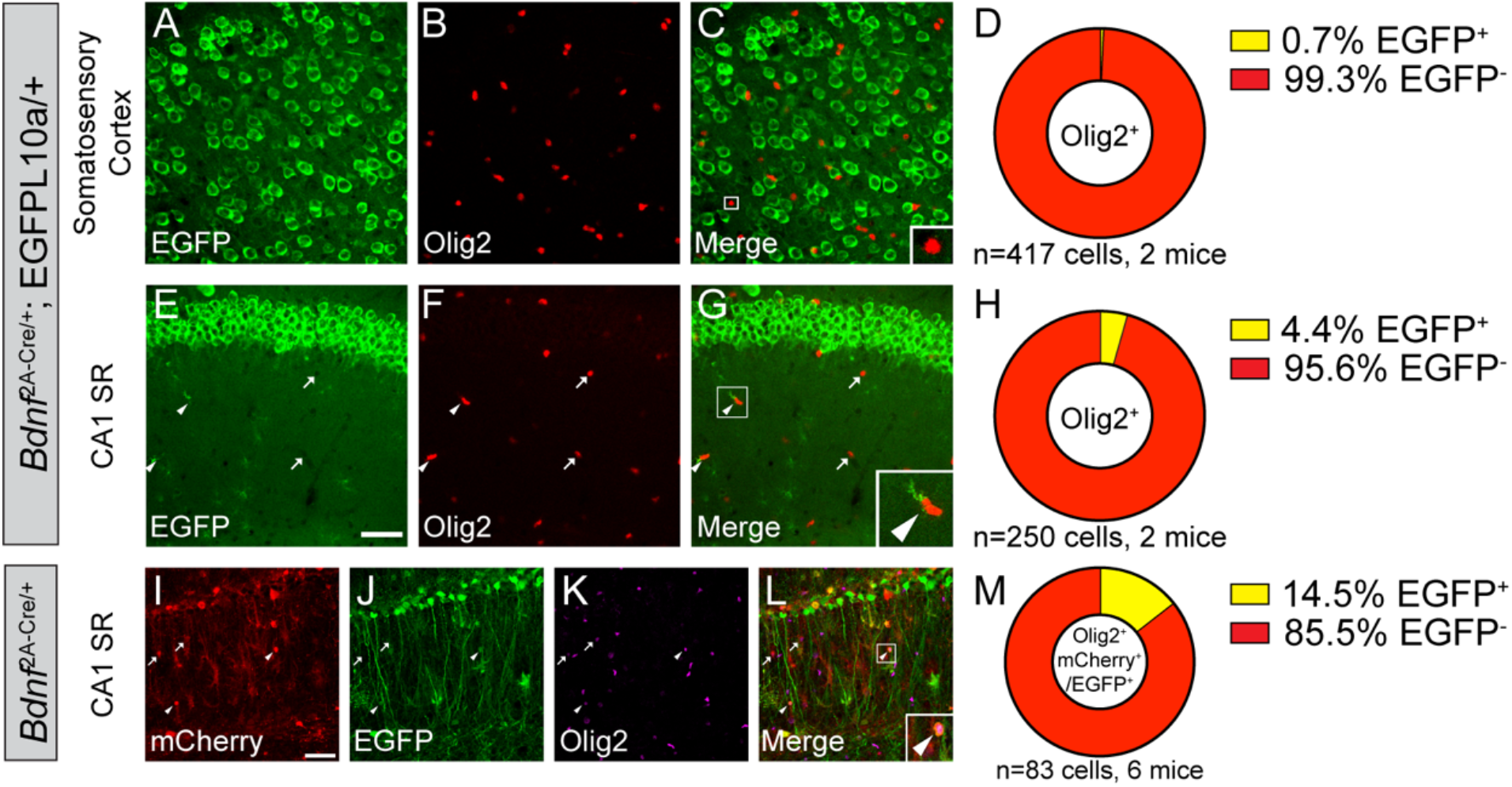
A small subset of oligodendrocytes express BDNF in the hippocampus. (**A-H**) Immunostaining of Olig2 in the SS (A-C) and CA1 SR (E-G) of adult *Bdnf*^2A-Cre/+^;EGFPL10a/+ mice, and (D,H) quantification of colocalization. (**I-M**) Immunostaining of Olig2 in the CA1 SR of adult *Bdnf*^2A-Cre/+^ mice injected with the mCherry-to-EGFP color switch AAV, and (M) quantification of colocalization. Scale bar = 50 μm. Arrowheads denote EGFP-expressing oligodendrocytes. Arrows denote oligodendrocytes that do not express EGFP.

### 3.4 Generation of *Ntrk2*^2A-Cre^ allele

To label cells that express TrkB.FL, we generated the *Ntrk2^2A-Cre/+^*mouse strain which express Cre recombinase in TrkB.FL-expressing cells. We employed the CRISPR/Cas9 technology to insert the P2A-Cre sequence into the *Ntrk2* locus immediately before the stop codon for the TrkB.FL (Fig. 5A). Mice with the insertion were identified using genomic DNA PCR (Fig. 5B). There was a high colocalization between *Cre* mRNA and *Ntrk2* mRNA for TrkB-FL in the cortex of *Ntrk2^2A-Cre/+^* mice (Fig. 5C-D), validating the allele.

**Figure 5.**
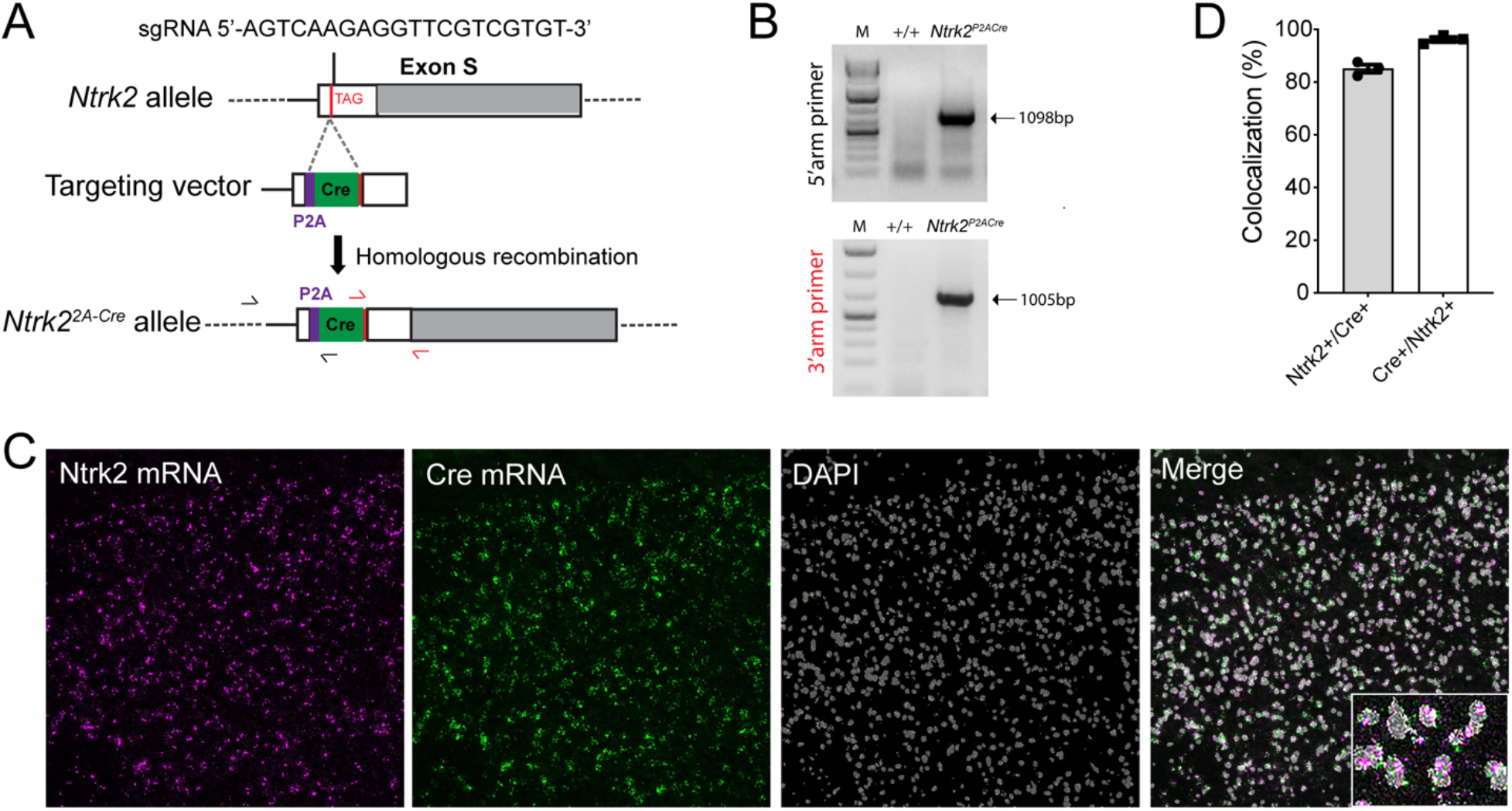
Generation of *Ntrk2^2A-Cre^* mice. (**A**) Strategy for insertion of the P2A-Cre sequence into the *Ntrk2* locus. (**B**) Genomic DNA PCR to identify *Ntrk2^2A-Cre/+^*mice. One primer is located outside each of the two homology arms in the targeting vector. (**C, D**) In situ hybridization of the cortex.

### 3.5 Microglia in the cortex, hippocampus or spinal cord do not express TrkB.FL or TrkB.T

To study TrkB.FL expression in glial cells, we crossed *Ntrk2*^2A-Cre/+^ mice that express Cre in TrkB.FL-expressing cells to the Cre-dependent EGFP-L10a/+ reporter mice to generate *Ntrk2*^2A-Cre/+^;EGFP-L10a/+ mice in which TrkB.FL expression is genetically labeled by EGFP. As expected, all neurons in the MO and hippocampal CA1 express TrkB.FL (Fig. 6A-F). Next, we examined whether microglia express TrkB.FL. No colocalization of EGFP and Iba1 was found in the MO or hippocampal CA1 SR of 4-month-old *Ntrk2*^2A-Cre/+^;EGFP-L10a/+ mice, indicating that microglia in these regions do not express TrkB.FL any time up to the time when the mice were perfused (Fig. 6G-L).

**Figure 6.**
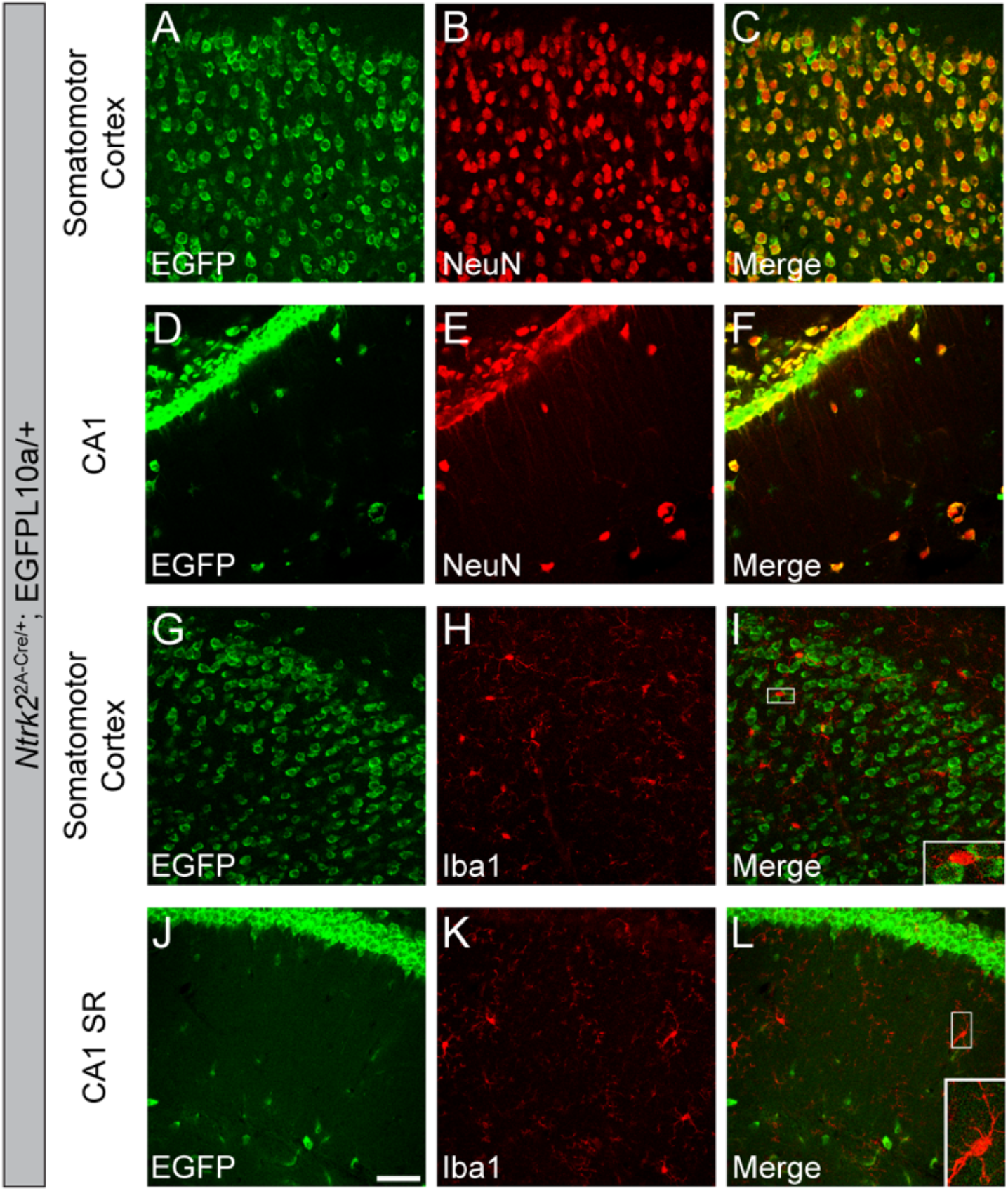
Microglia do not express TrkB.FL. (**A-F**) Colocalization of NeuN^+^ neurons and EGFP in the MO and CA1 of *Ntrk2*^2A-Cre/+^;EGFP-L10a/+ mice. (**G-L**) Homeostatic microglia do not express EGFP in the MO and CA1 SR of *Ntrk2*^2A-Cre/+^;EGFP-L10a/+ mice. Scale bar = 50 μm. n = 2 mice.

To see whether microglia express TrkB.T, we utilized *Ntrk2*^CreER/+^ mice that express Cre-ER under the *Ntrk2* promoter. We crossed *Ntrk2*^CreER/+^ mice to Cre-dependent EGFP-L10a/+ reporter mice to generate *Ntrk2*^CreER/+^;EGFP-L10a/+ mice, in which cells expressing either TrkB.FL or TrkB.T are genetically labeled by EGFP upon tamoxifen exposure. As expected, neurons in the cortex and hippocampal CA1 are labeled by EGFP due to their expression of TrkB.FL and TrkB.T. However, only a subset of neurons were labeled by EGFP (Fig. 7A-F). Because almost all neurons should express TrkB.FL in adult mice (Fig. 6A-F), this suggests that the Cre induction by tamoxifen was lower than 100%. Our data showed no colocalization of EGFP and Iba1 in either MO or CA1 SR region (Fig. 7G-L). This means that Iba1^+^ microglia do not express either TrkB.FL or TrkB.T in adult mice.

**Figure 7.**
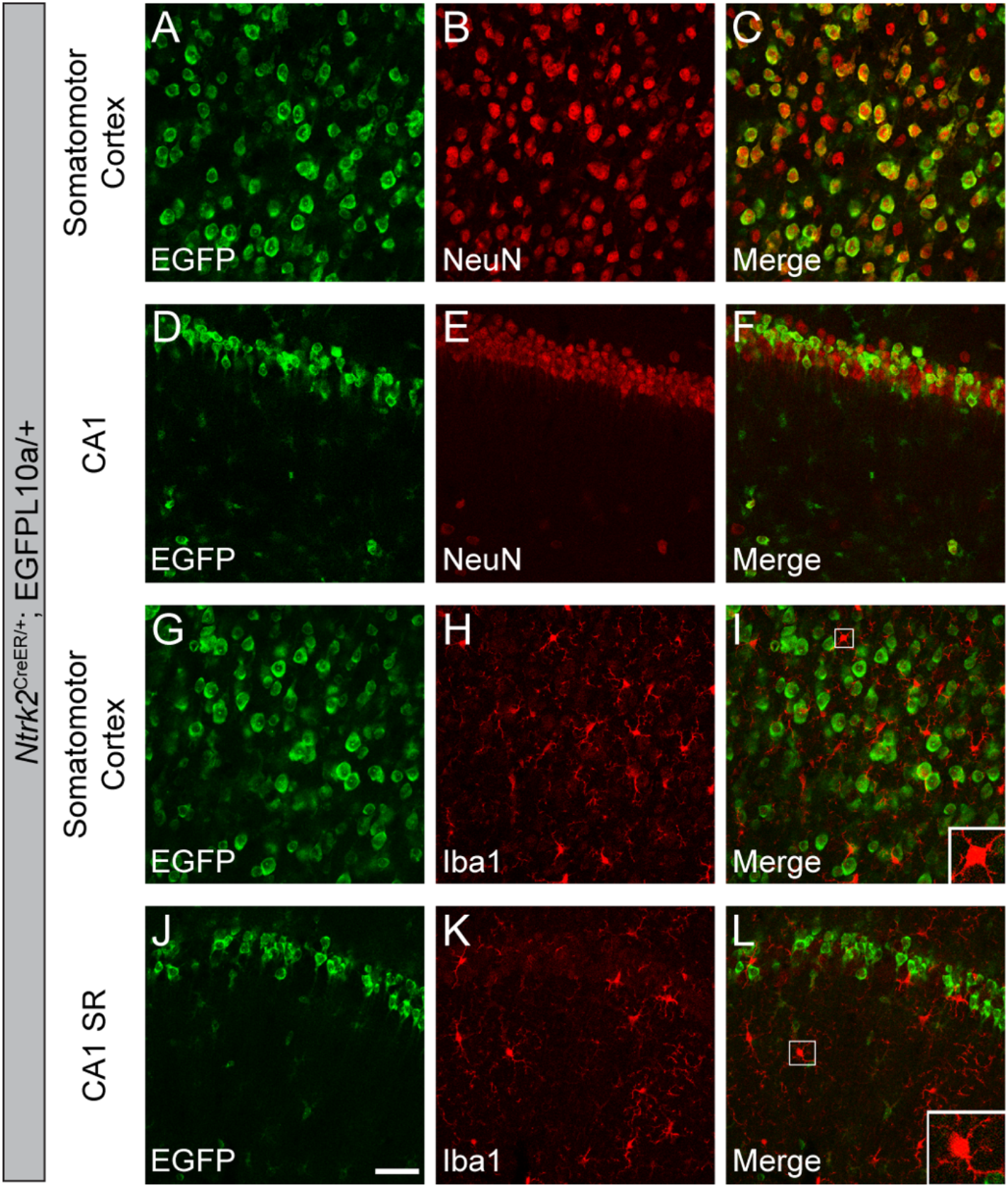
Microglia do not express either TrkB.FL or TrkB.T in adult mice. (**A-F**) Colocalization of NeuN^+^ neurons and EGFP in the MO and CA1 of *Ntrk2*^CreER/+^;EGFP-L10a/+ mice. (**G-L**) Homeostatic microglia are not labeled by EGFP in the MO and CA1 SR of *Ntrk2*^CreER/+^;EGFP-L10a/+ mice. Scale bar = 50 μm. n = 2 mice.

### 3.5 Astrocytes predominantly express TrkB.T and a small subset of astrocytes express TrkB.FL

Astrocytes have been shown to predominantly express TrkB.T; however, whether astrocytes also express TrkB.FL is not as clear. To investigate the expression of TrkB.FL in astrocytes, we quantified colocalization of EGFP and astrocyte markers, SOX9 and GFAP, in *Ntrk2*^2A-Cre/+^;EGFP-L10a/+ mice. Our analyses showed that a small portion of astrocytes (16.8% in SS and 18.0% in hippocampal CA1 SR) express TrkB.FL at some point during the lifespan of the mice (Fig. 8A-H). To test whether astrocytes in adult mice express TrkB.FL, we injected the Cre-dependent mCherry-to-EGFP color switch AAV to the hippocampus of 2-month-old *Ntrk2*^2A-Cre/+^ mice [24]. 92.7% of astrocytes were transduced by the AAV virus, and only 3.6% of the transduced cells express Cre (Fig. 8I-M). These results suggest that only a very small subset of astrocytes express TrkB.FL in adult mice. Because this number is smaller than the 18.0% seen in *Ntrk2*^2A-Cre/+^;EGFP-L10a/+ mice. The two possibilities discussed before can also apply here: (1) A small subset of astrocytes express TrkB.FL only early during development and stop the expression in adult mice; (2) at each different time point, there is a different small subset of astrocytes that express TrkB.FL, and that accumulates to the 18% seen in *Ntrk2*^2A-Cre/+^;EGFP-L10a/+ mice.

**Figure 8.**
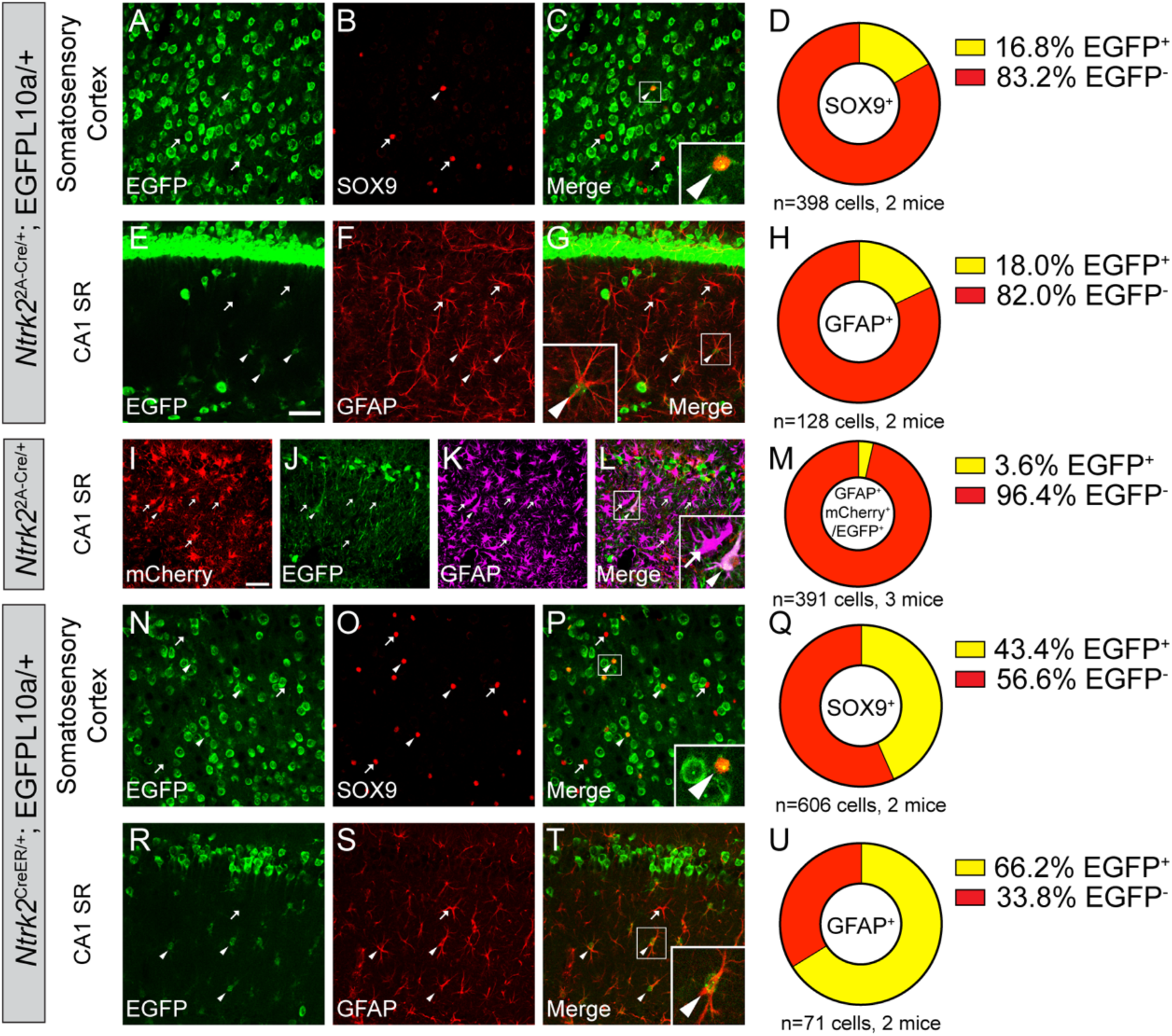
Astrocytes predominantly express TrkB.T in adult mice. (**A-G**) Immunostaining of SOX9 (A-C) or GFAP (D-G) in the SS (A-C) and CA1 SR (E-G) of *Ntrk2*^2A-Cre/+^;EGFP-L10a/+ mice, and (D,H) quantification of colocalization. (**I-M**) Immunostaining of GFAP in the CA1 SR of adult *Ntrk2*^2A-Cre/+^ mice injected with the mCherry-to-EGFP color switch AAV, and the quantification of the colocalization of GFAP and EGFP. (**N-U**) Immunostaining of SOX9 (A-C) or GFAP (D-G) in the SS (A-C) and CA1 SR (E-G) of *Ntrk2*^CreER/+^;EGFP-L10a/+ mice, and the quantification of colocalization. Scale bar = 50 μm. Arrowheads denote EGFP-expressing astrocytes. Arrows denote astrocytes that do not express EGFP.

Next, we examined expression of TrkB.T using *Ntrk2*^CreER/+^;EGFP-L10a/+ mice where cells expressing either TRKB.FL or TrkB.T are labeled by EGFP. Tamoxifen was given at the age of 6 weeks to induce Cre recombination. We found that 43.4% astrocytes express either TrkB.FL or TrkB.T in the SS, and 66.2% in the CA1 SR (Fig. 8N-U). Because we showed that most astrocytes do not express TrkB.FL in the CA1 SR of adult mice (Fig. 8I-M), this means most of the EGFP^+^ astrocytes in the *Ntrk2*^CreER/+^;EGFP-L10a/+ mice express TrkB.T only. Due to the incomplete Cre induction described above, the actual percentage of astrocytes that express TrkB in *Ntrk2*^CreER/+^;EGFP-L10a/+ mice could be higher than what we show.

### 3.6 Oligodendrocytes express TrkB.FL and TrkB.T in adult brains

Lastly, we investigated TrkB expression in oligodendrocytes. Colocalization of oligodendrocyte marker Olig2 and EGFP in *Ntrk2*^2A-Cre/+^;EGFP-L10a/+ mice suggests that a subset of oligodendrocytes (15.8% in SS and 43.4% in CA1 SR) express TrkB.FL at some point during their lifespan (Fig. 9A-H). To examine whether oligodendrocytes express TrkB.FL in adult mice, the Cre-dependent mCherry-to-EGFP color switch AAV was injected to the hippocampus of adult *Ntrk2*^2A-Cre/+^ mice. The transduction rate of oligodendrocytes is lower than that of astrocytes. Around 27.8% of oligodendrocytes were transduced by the AAV. We observed that 21.1% of transduced Olig2^+^ oligodendrocytes express TrkB.FL in the adult mouse brains (Fig. 9I-M).

**Figure 9.**
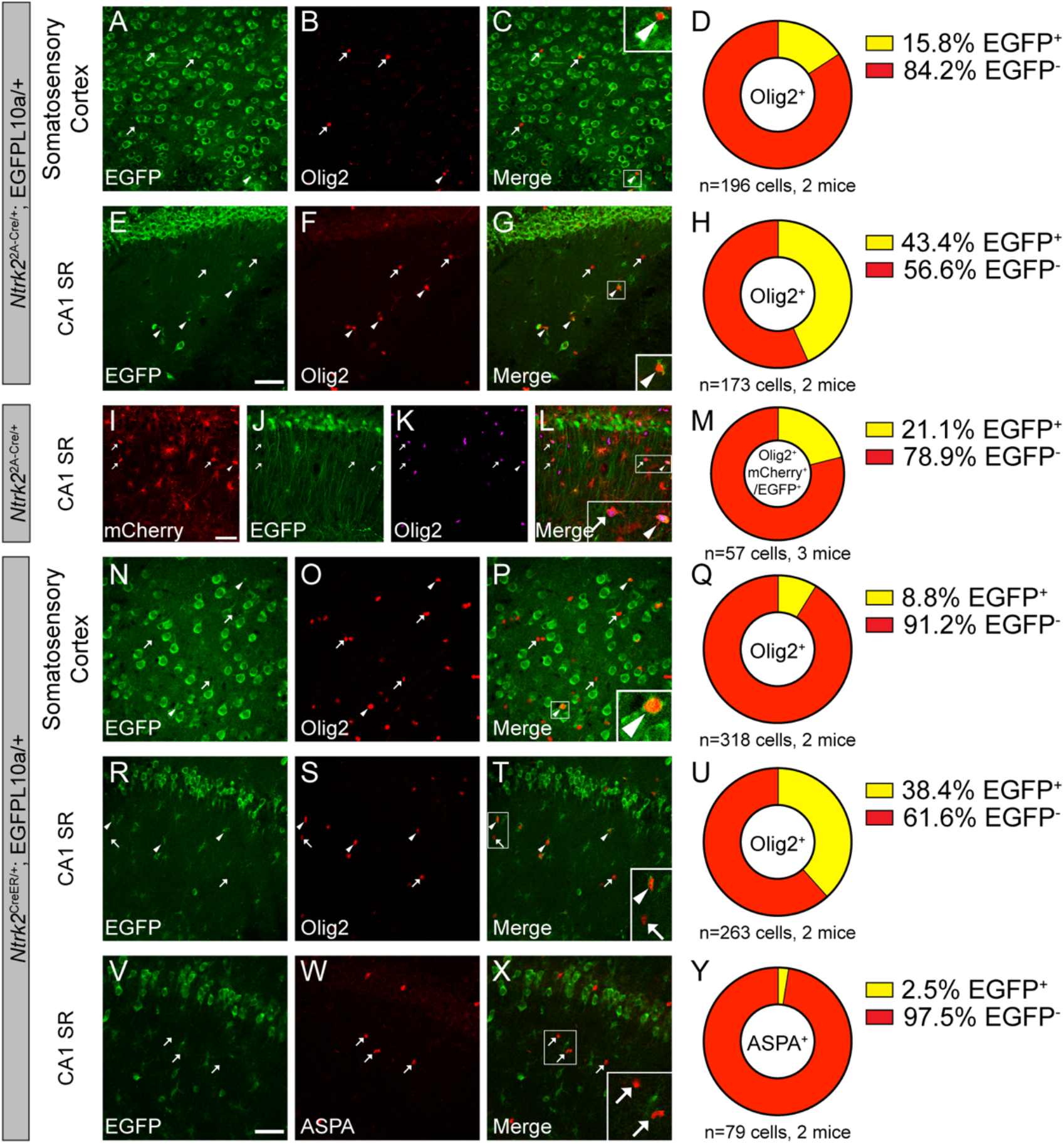
OPCs express TrkB.FL and TrkB.T in adult mice. (**A-G**) Immunostaining of Olig2 in the SS (A-C) and CA1 SR (E-G) of *Ntrk2*^2A-Cre/+^;EGFP-L10a/+ mice, and (D,H) quantification of colocalization. (**I-M**) Immunostaining of Olig2 in the CA1 SR of adult *Ntrk2*^2A-Cre/+^ mice injected with the mCherry-to-EGFP color switch AAV, and quantification of the colocalization of GFAP and EGFP. (**N-U**) Immunostaining of Olig2 in the SS (A-C) and CA1 SR (E-G) of *Ntrk2*^CreER/+^;EGFP-L10a/+ mice, and (Q,U) quantification of colocalization. (**V-Y**) Immunostaining of ASPA in the CA1 SR (of *Ntrk2*^CreER/+^;EGFP-L10a/+ mice, and (Y) quantification of colocalization. Scale bar = 50 μm. Arrowheads denote EGFP-expressing oligodendrocytes. Arrows denote oligodendrocytes that do not express EGFP.

To find out whether oligodendrocytes also express TrkB.T, we quantified colocalization of Olig2 and EGFP in tamoxifen-treated *Ntrk2*^CreER/+^;GFPL10/+ mice. We found that 8.8% of Olig2^+^ oligodendrocytes are colocalized with EGFP in the SS, indicating that these cells express either TrkB.FL or TrkB.T in the adult mice. In the CA1 SR, 38.4% of Olig2^+^ oligodendrocytes express either TrkB.FL or TrkB.T in the adult mice (Fig. 9R-U). Because we showed 21.1% of Olig2^+^ oligodendrocytes express TrkB.FL in the CA1 SR of adult mice, this means that there are at least another 17% of Olig2^+^ oligodendrocytes that express TrkB.T in the adult mouse brains. Olig2 labels both oligodendrocyte precursor cells (OPCs) and mature oligodendrocytes. To differentiate between these two populations, we used aspartoacylase (ASPA) as a marker to mark mature oligodendrocytes only. Our results show that nearly all mature oligodendrocytes do not express either TrkB.FL or TrkB.T in the CA1 SR of adult mice, which suggests that only OPCs express TrkB.FL and TrkB.T in adult mice (Fig. 9V-Y).

## 4. Discussion

Whether microglia express BDNF and TrkB or not has been debated. Microglia were discovered to express BDNF *in vitro* at the level of mRNA [26, 27] and protein [28] decades ago. LPS stimulation was suggested to increase BDNF secretion from cultured microglia [26, 28]. *In vivo* experiments showed BDNF expression in activated microglia [29]. BDNF from spinal microglia were suggested to be responsible for pain hypersensitivity [7]. Microglial BDNF was also reported to promote motor learning-dependent synapse formation [6]. Following these studies, many researchers showed the knockdown of microglial expression of BDNF disrupts biological processes including self-renew and proliferation of hippocampal neurons [30], nerve injury-induced pyramidal neuron hypersensitivity in the SS cortex [31], and mechanical allodynia induced by high-frequency stimulation [32]. Microglial BDNF was also suggested to be critical for (*R*)-ketamine-mediated microglial activation in the medial prefrontal cortex of chronic social defeat stress mice [33]. However, other studies using *Bdnf^Laz/+^* reporter mice and transcriptomic analysis show that microglia do not express significant level of BDNF in the spinal cord [8, 34]. This inconsistency likely comes from the lack of binding specificity of BDNF antibodies, as many previous efforts to characterize BDNF expression relied on immunohistochemistry. To avoid the use of BDNF antibodies and increase sensitivity, we took advantage of the *Bdnf*^2A-cre/+^ knockin mice that express Cre recombinase under the *Bdnf* promoter and the EGFPL10a/+ reporter mice that express EGFP localized in cell bodies but not dendrites and axons for better characterization of colocalization. Our data suggest that both resting and activated microglia do not express BDNF in the MO cortex, hippocampal CA1, and spinal cord. These results are consistent with findings from transcriptomic analysis that showed microglia do not express BDNF in the spinal cord [8], and from another group that recently demonstrated resting microglia and microglia activated by ATP do not express BDNF transcriptionally and translationally in the mouse motor cortex [35]. Therefore, the phenotypes seen after deleting the microglial *Bdnf* gene likely come from artifacts. However, our data do not exclude the possibility that microglia may have the potential to internalize and recycle BDNF expressed by neurons and other cell types in the CNS.

Using similar method, we also show that microglia do not express the TrkB receptor in the brain and spinal cord. Our data show microglia do not express TrkB.FL from embryo to adult, and do not express TrkB.T in adult mice. This means that at least in adult mice, microglia do not respond to BDNF through the TrkB receptor. However, it is still likely that BDNF can signal through the low affinity receptor p75^NTR^ in microglia, as studies have suggested a subset of microglia express p75^NTR^ and its expression is upregulated upon infection in mice [36]. Another study suggested no significant p75^NTR^ expression in wild-type microglia, but the expression is increased in 5xFAD mice and Alzheimer’s patients [37], suggesting a potential role of BDNF-p75^NTR^ signaling in neuroinflammation.

Cultured astrocytes have been shown to express BDNF and the expression is increased by KCl, carbachol, and glutamate [38, 39]. Astrocytic BDNF regulates neuronal dendrite complexity and spine number *in vitro* [16]. Studies have suggested that cortical layer II/III astrocytes can incorporate extracellular proBDNF through p75^NTR^-mediated endocytosis and cleave proBDNF to mBDNF [15]. But whether and what percentage of astrocytes produce BDNF *in vivo* under physiological condition remain elusive. Our results show that a small subset of astrocytes express BDNF in the SS cortex (14.6%) and hippocampal CA1 SR (18.3%) under physiological condition in mice. In adult mice, 10% of astrocytes in CA1 SR express BDNF. This is consistent with other studies that showed a relatively low baseline level of mBDNF in astrocytes [40], but an upregulation of astrocytic BDNF expression under pathological conditions induced by MK-801, an N-methyl-D-aspartic acid (NMDA) receptor antagonist, *in vitro* [41] and treatment of glatiramer acetate in R6/2 HD mouse model *in vivo* [40]. Although only 10% of astrocytes express BDNF in the hippocampus under physiological condition, astrocytes seem to be an important source of BDNF for normal CNS functions. Studies using BDNF knock-down mouse models found that astrocytic BDNF is required for oligodendrogenesis after white matter damage [42]. In the experimental autoimmune encephalomyelitis (EAE) model of multiple sclerosis, astrocyte specific BDNF knockout led to more severe clinical course with increased axonal injury and loss [43].

It is known that astrocytes predominantly express TrkB.T, which plays important roles in mediating calcium release from intracellular stores in astrocytes [9] and controlling astrocyte morphological maturation [11]. In the adult brain, only less than 5% astrocytes express TrkB.FL in the hippocampal CA1 SR. These data are consistent with the current belief that astrocytes dominantly express TrkB.T in the adult mice. We also show that cumulatively through embryo to adult, 18% astrocytes in the hippocampal CA1 SR have expressed TrkB.FL, suggesting it is likely that some astrocytes only express TrkB.FL early during embryonic and postnatal developmental stages. This corroborates a previous *in vitro* study that showed radial glia (the main astrocytic precursor cells) and proliferating cortical astrocytes in culture express mRNAs for both TrkB.FL and TrkB.T, whereas differentiating astrocytes predominantly express TrkB.T and downregulate TrkB.FL expression [44]. These data suggest that BDNF-TrkB.FL signaling may play a role in astrocyte development and maturation.

Our data show that a very small subset (less than 5%) of oligodendrocytes in the hippocampal CA1 SR express BDNF. Contradictory to the studies that showed BDNF expression in cortical oligodendrocytes [45, 46], our data showed that most Olig2^+^ cells in the cortex do not express BDNF. Oligodendroglial BDNF expression has been reported by a few studies. A study has showed that oligodendrocytes in the spinal cord and optic nerve express BDNF mRNA and the expression is increased in Long-Evans shaker rats that lack CNS myelin due to the MBP gene mutation [47]. Another study found that fingolimod treatment resulted in increased BDNF mRNA concomitant with increased OPC differentiation in rat OPC culture [48]. Our results confirm the expression of BDNF in oligodendrocytes in certain brain regions. And our observation of increased BDNF-expressing oligodendrocytes in AAV injected mice may suggest that BDNF expression is increased when myelin is disrupted, as a previous study has reported increased BDNF level in oligodendrocytes after injury [49].

Our results also showed that only OPCs, but not mature oligodendrocytes, express TrkB.FL and TrkB.T in the hippocampal CA1 SR of adult mice. BDNF-TrkB signaling in OPCs has been suggested to have important functions. OPC-specific TrkB deletion resulted in impaired activity-dependent myelination [18]. BDNF and its structural mimetic TDP6 have been shown to promote myelination and myelin repair via oligodendroglial TrkB receptor [20, 50, 51]. Future studies are necessary to determine which TrkB isoform(s) are responsible for mediating these effects.

In summary, we characterized the expression of BDNF and TrkB in microglia, astrocytes, and oligodendrocytes in the current study. Our results clarify that microglia in the examined regions do not express BDNF or TrkB. Our findings also open opportunities to investigate previously unidentified roles of BDNF and TrkB in astrocytes and oligodendrocytes.

## Author Contributions

C.N. and B.X. designed research; C.N., X.Y., J.J.A., and H.X. performed experiments; C.N. analyzed data; C.N. wrote and B.X. edited the paper.

## Data Availability Statement

Original data generated and analyzed during this study are included in this published article.

## Acknowledgments

We thank David Ginty for the *Ntrk2^CreER^*mouse strain and Zhonghou Wang for injecting viruses into the hippocampus in a few mice.

